# Skeletal muscles that actuate sexual displays are specialized for *de novo* androgen synthesis

**DOI:** 10.1101/740720

**Authors:** Eric R. Schuppe, Franz Goller, Matthew J. Fuxjager

## Abstract

The gonads (testes) act as the primary organ where androgenic hormones are made to regulate reproductive behavior in male vertebrates. Yet many endocrinologists have also long suspected that other tissues in the body can autonomously synthesize their own androgens to support behavioral output. We examine this idea here by studying whether avian skeletal muscles that actuate elaborate socio-sexual displays are specialized to maintain the molecular machinery otherwise needed for de novo androgen synthesis. Our results show that the vocal organ, or syrinx (SYR), of two songbirds species does in fact express all transporters and enzymes necessary to create androgenic hormones from scratch. This includes genes that encode proteins to mediate rate-limiting steps of steroidogenesis, which are seldom found outside of the gonads. We also show that expression levels of these genes are far greater in the SYR than non-display muscles, matching expression levels found in another extra-gonadal site of steroidogenesis—the brain. Furthermore, we uncover a nearly identical gene expression signature in a woodpecker neck muscle, the *longus colli ventralis* (LC). This tissue actuates the bird’s social drum display, which serves the same functions as song. This same study also demonstrates that the elevated expression of steroidogenic genes persists in this neck muscle year-round, suggesting that the LC’s capacity to make androgens is a constitutive trait. Altogether, our findings suggest that muscles involved in sexual display is uniquely specialized to locally make steroid hormones, likely supporting their own role in behavioral production.

## INTRODUCTION

Androgenic hormones play a vital role in the regulation of male reproductive behavior, including most forms of courtship and male-male competition [1]. Conventional wisdom says that the gonads (testes) mediate these effects, given that they synthesize androgenic steroids *de novo* and release them into circulation [2, 3]. Yet, the dependency of reproductive behavior on the gonads may be more complex than this model suggests. Many species, for example, express androgen-dependent behavioral traits at times when the testes are fully regressed and incapable of producing steroids [4-8]. Such results imply that androgen synthesis may occur in other tissues throughout the vertebrate body, locally fueling steroidal modulation of behavioral output. Although this idea provides an intriguing way to conceptualize how behavior can be adaptively decoupled from testicular functioning [9, 10], our understanding of where extra-gonadal androgen synthesis occurs and how it is specialized to support sexual traits remains largely unclear.

Here we explore these topics by assessing mechanisms of *de novo* androgen synthesis in skeletal muscles that support elaborate socio-sexual behavior. Skeletal muscle is a major androgen target in the vertebrate body [11-13], and prior work shows that androgenic regulation of muscle is often necessary for reproduction [14-16]. Courtship is a prime example: muscles that mediate displays used to attract mates are often enriched with androgen receptor [13, 17, 18], and blocking androgenic activation of these receptors disrupts the effectiveness or recognizability of signaling behavior [14-16]. Importantly, however, we know that individuals can often produce these same displays when their gonads are not making steroids [4, 5, 19]. Tissues other than the testes may therefore act as the source of androgens to regulate how muscle governs behavioral outflow. One potential site of androgen synthesis is the display muscle itself—that is, myocytes may make their own androgens to locally influence their ability to control behavior. Yet we know little about the capacity of skeletal muscles to create its own androgens, making this hypothesis difficult to assess.

One of the best ways to examine whether a given tissue can make its own steroid hormones *de novo* is to measure how it expresses the transporters and enzymes that make up the steroid biosynthesis pathway [Figure 1; 20, 21, 22]. Indeed, there are only a few tissues other than the gonads that readily express these genes—namely, skin, heart, and brain [22-24]. Most other tissues in vertebrates lack key transporters and enzymes that mediate steroid biosynthesis, particularly the rate-limiting step of this process where cholesterol is brought into the mitochondria and converted to the steroid hormone precursor, pregnanolone [25-27]. Accordingly, if selection drives the evolution of *de novo* androgen synthesis in displays muscles, then these tissues should express the rate-limiting transporters and enzymes that facilitate this important first steroid of steroidogenesis as well as the remaining enzymes that metabolize pregnanolone into testosterone. Moreover, if muscles are specialized to do this, then we also expect that they express the genes more abundantly than muscles not involved in display production. In fact, such expression should be on par with other extra-gonadal sites of *de novo* steroid synthesis, like the brain [22, 28].

**Figure 1.**
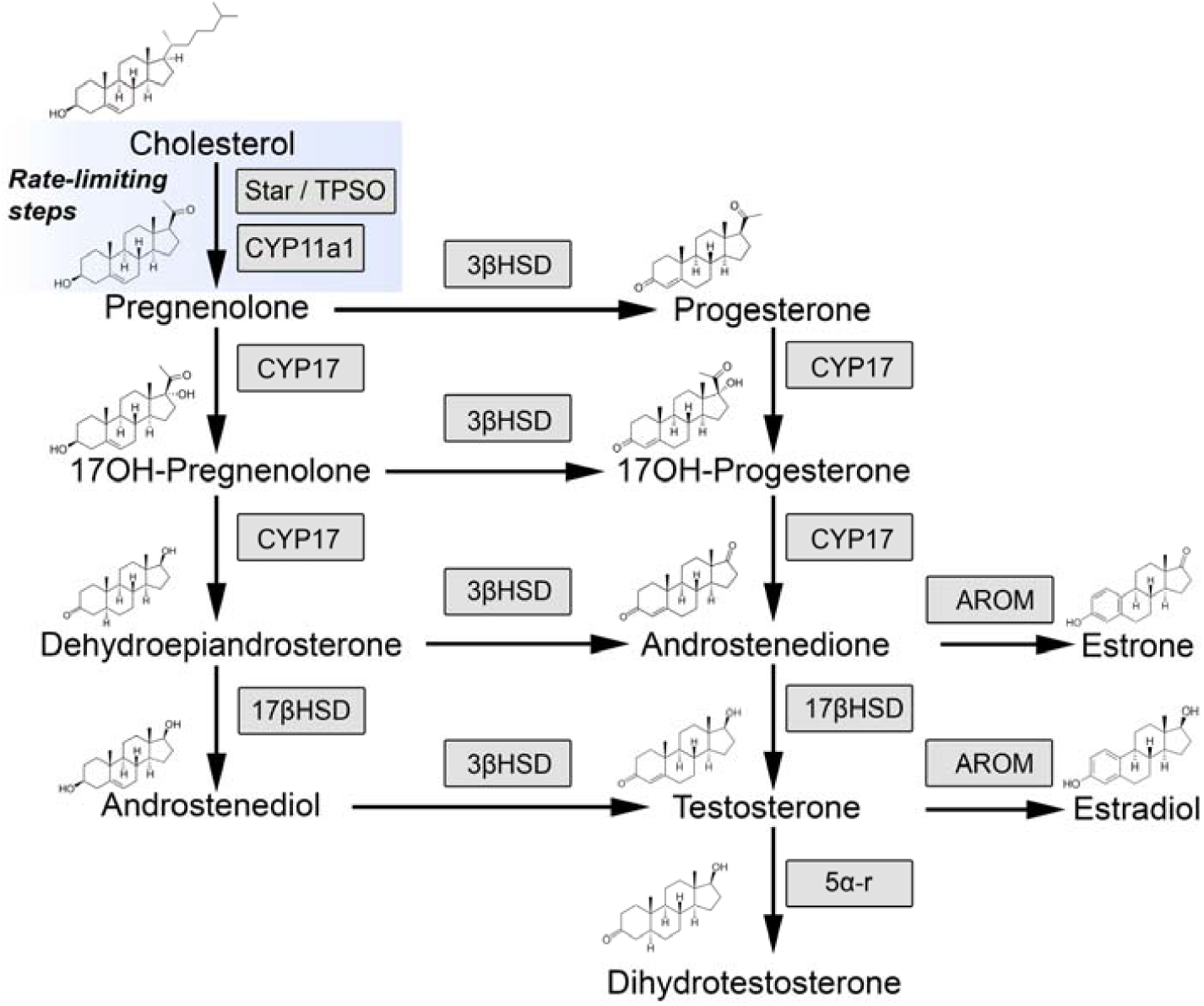
Pathway that outlines the multi-step conversion of cholesterol to androgenic (red box) and estrogenic (green box) hormones. The rate limiting proteins in *de novo* steroid synthesis (blue box) play an important role in the initial transport (steroidogenic acute regulator protein [StAR] and translocator protein [TPSO]) and the subsequent enzymatic conversion (cyp11a1) of cholesterol to pregnenolone. Other enzymes responsible for the conversion of this hormone to other steroid metabolites, including androgens (17β-hydroxysteroid dehydrogenase [17BHSD] and 3-oxo-5α-steroid 4-dehydrogenases [5α-r]) and estrogens (Aromatase [AROM]). The latter (AROM) is also important because it act through intracellular estrogen receptor, which are not specialized in muscles that participate in sexual signaling [18]. All enzymes are highlighted in grey rectangles.

We test these predictions in three different bird species: white-breasted nuthatches (*Sitta carolinensis*), zebra finches (*Taeniopygia guttata*), and downy woodpeckers (*Dryobates pubescens*). The first reason that these taxa are ideal to study muscular androgen synthesis is that they each produce an elaborate socio-sexual display that depends on androgen-muscle interactions. Nuthatches and zebra finches, for example, court mates and compete with rivals by singing [29, 30]. This behavior is controlled by a vocal organ called the syrinx (SYR), which is composed of several muscles that sit on the tracheo-bronchial junction [31]. Neural activation of this musculature shapes the spectral features of song [32], and studies show that androgenic hormones fine tune this process by acting directly on androgen receptors in SYR muscle [11, 15]. On the other hand, downy woodpeckers signal to conspecifics during territorial contests by drumming their bill on trees at speeds averaging 16 beats/sec [33]. Such behavior is mediated by a neck muscle called the *longus colli ventralis* (LC) [34], and recent work shows that androgens likely act on it to help support the performance of drumming behavior [35]. The second reason why these three species are ideal for the proposed work is that they all are known to produce their respective displays when circulating androgens levels are basal, including periods of non-breeding [5, 30, 36]. We therefore test whether the SYR and LC muscles of these birds not only express the necessary machinery for *de novo* androgen synthesis, but also whether these tissues are specialized for such endocrine functionality.

## RESULTS

### Muscular expression of steroid biosynthesis enzymes

We first tested whether display muscles express the necessary transporters and enzymes for *de novo* androgen synthesis. Using reverse transcription PCR, we assayed the presence of transcripts for these proteins in both the white-breasted nuthatch SYR and downy woodpecker LC. Our data showed that all these transcripts were present in both display muscles, although some were expressed at very low levels (Figure S1). We also included testes in these PCR reactions as a positive control, verifying our primers and their efficacy in the PCR reactions. Overall, our data indicate that all transporters and enzymes needed to turn cholesterol into an androgenic ligand (see Figure 1) are present in both the nuthatch SYR and the woodpecker LC.

### *Specializations in the capacity for* de novo *androgen synthesis in the syrinx*

Next, we explored whether the machinery for *de novo* androgen synthesis is specialized in display muscle. To do this, we used real-time quantitative PCR (qPCR) to compare relative expression of androgen biosynthesis genes among the (*i*) gonad (testis), which are the main site of androgen synthesis; (*ii*) brain, which is a known site of extra-gonadal androgen synthesis; (*iii*) SYR and/or LC display muscles, and (*iv*) *pectoralis* muscle (PEC), a large non-display muscle that powers locomotion in these species.

In both nuthatches and zebra finches, we found significant differences among tissues in the expression of nearly all androgen biosynthesis genes (Figure VI-3A and B; Table VI-1). The one exception was zebra finch TPSO, which was highly expressed in every tissue. *Post-hoc* analyses of these main effects showed that gonads have the highest expression of each gene. At the same time, our findings also highlighted that SYR muscle expresses these genes at similar levels to the brain, another extragonadal site of *de novo* steroid synthesis [22]. Gene expression levels in SYR muscle were also significantly greater than in a non-display muscle, the PEC (Figure VI-3A and B; Table VI-1). Only cytochrome p450 17 (cyp17, or steroid 17α-monooxygenase) showed an expression profile in SYR that was lower than in the brain and equal to the PEC. Cyp17 converts progestins to androgens [37], its abundance is unlikely to limit the tissue’s capacity for androgen synthesis.

Perhaps the most important result of this analysis is that expression patterning described above applies to both steroidogenic acute regulatory protein (StAR) and cytochrome p450 family 11 (cyp11a1, or side chain cleavage enzyme). These gene products mediate the rate-limiting steps of steroidogenesis—StAR transports cholesterol from the outer mitochondrial membrane to the inner membrane [38], while side chain cleavage enzymatically converts cholesterol to pregnanolone [37]. Most extra-gonadal tissues fail to express either of these genes, rendering them incapable of *de novo* steroid synthesis [25-27]. Thus, display muscle is unique in this regard, as it expresses relatively high levels of these genes to likely initiate the process of steroidogenesis.

One of the marked species differences in the SYR is that the nuthatch expresses aromatase, while the zebra finch does not (Figure 2A and B; Table 1). Aromatase converts the androgen testosterone to estradiol, meaning that these species likely differ in the amount of estrogens that the SYR can make. To date, most work suggests that estrogens play little role in the regulation of the SYR; yet, these results suggest that this may not be the case for all birds. Notably, both the nuthatch and zebra finch SYR express 5α-reductase, which converts testosterone to a more potent androgenic ligand (dihydrotestosterone).

**Table 1.** Summary of linear mixed models used to assess tissue differences in steroidogenic enzyme expression in male zebra finches, white-breasted nuthatches, and downy woodpeckers

|  | **STAR** | **TPSO** | **Cyp11a1** | **3BHSD** | **Cyp17** | **17BHSD** | **5aR** | **AROM** |
| --- | --- | --- | --- | --- | --- | --- | --- | --- |
| **Zebra finch** | F<sub>3,10</sub>=62.40,<br>p<0.001 | F<sub>3,10</sub>=3.32,<br>p=0.07 | F<sub>3,10</sub>=8.64,<br>p<0.01 | F<sub>3,10</sub>=74.84,<br>p<0.001 | F<sub>3,10</sub>=31.18,<br>p<0.001 | F<sub>3,10</sub>=8.34,<br>p<0.01 | F<sub>3,10</sub>=17.14,<br>p<0.001 | F<sub>3,10</sub>=59.85,<br>p<0.001 |
| **White-breasted nuthatch** | F<sub>4,12</sub>=22.64,<br>p<0.001 | F<sub>4,10</sub>=26.79,<br>p<0.001 | F<sub>4,11</sub>=27.07,<br>p<0.001 | F<sub>4,11</sub>=48.23,<br>p<0.001 | F<sub>4,11</sub>=21.03,<br>p<0.001 | F<sub>4,11</sub>=68.12,<br>p<0.001 | F<sub>4,12</sub>=26.30,<br>p<0.001 | F<sub>4,12</sub>=33.04,<br>p<0.001 |
| **Downy woodpecker** | F<sub>3,10</sub>=33.62,<br>p<0.001 | F<sub>3,7.13</sub>=27.89,<br>p<0.001 | F<sub>3,10</sub>=7.33,<br>p<0.01 | F<sub>3,7.57</sub>=217.71,<br>p<0.001 | F<sub>3,10</sub>=58.01,<br>p<0.001 | F<sub>3,7</sub>=80.77,<br>p<0.001 | F<sub>3,7.88</sub>= 104.66,<br>p<0.001 | F<sub>3,10</sub>=22.38,<br>p<0.001 |

\* denote significant differences after FDR correction for multiple tests

**Figure 2.**
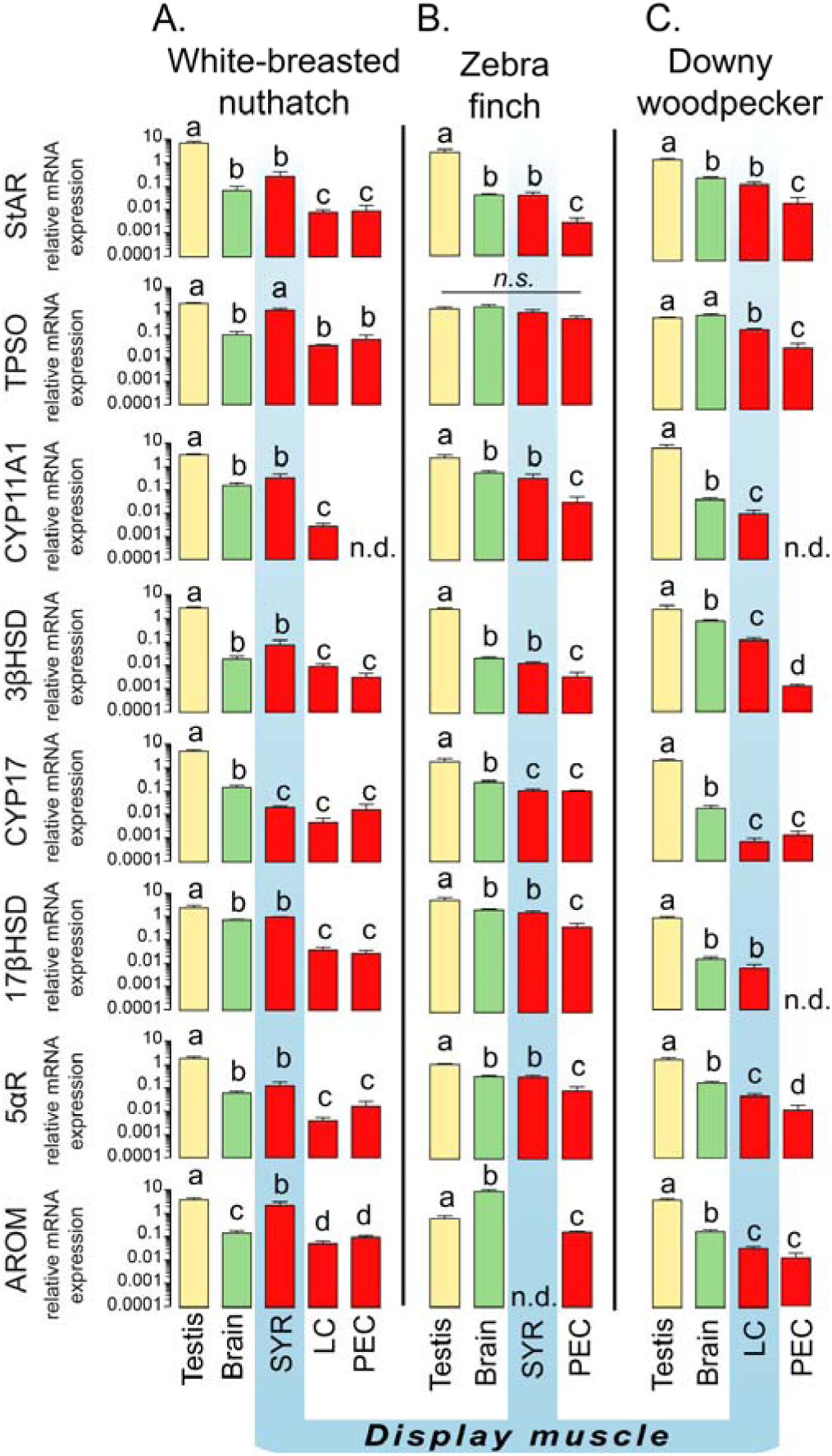
Expression of steroidogenic enzymes in the testis (yellow), brain (green) and skeletal muscle (red) of the downy woodpecker, zebra finch, and white-breasted nuthatch. Display muscles (e.g. the *longus colli ventralis* (LC) and *syrinx* (SYR) are highlighted with a blue line. For all species, the non-display muscle was the *pectoralis* (PEC). Bars represent mean relative mRNA expression (gene interest/GAPDH) ±SEM. Letters denote *post hoc* significant differences (p <0.05) between tissues. Non-detectable transcripts are denoted as ‘nd,’ indicating a failure to detect any amplification after 40 cycles. The only exception to this was expression of cyp11a1 listed as n.d. for the white-breasted nuthatch PEC. The Ct was ≈ 37, which is greater than threshold we used to determine functional “detectability” with confidence [76].

### Steroid biosynthesis in the woodpecker neck muscle

In a second experiment, we explored the expression profile of these same androgen biosynthesis genes in the downy woodpecker neck muscle, the LC [39]. This tissue actuates the birds drum display, which is functionally similar to birdsong but produced through hammering the bill against a tree (and not via the SYR). We find that LC muscle expresses all transporters and enzymes necessary for androgen biosynthesis, including the transcripts for both StAR and side chain cleavage (Figure 2C; Table 1). Moreover, each gene is more abundantly expressed in the LC than in the PEC (non-display muscle). The one exception to this pattern is cyp17, whose expression levels are indistinguishable between the two muscles (Figure 2C; Table 1). Importantly, this difference between the LC and PEC may be associated specifically with drumming in woodpeckers, as no such difference in gene expression profile was present between these two muscles in the nuthatch.

Meanwhile, in the downy woodpecker, transcript levels are typically lower in the LC and PEC, compared to both the brain and gonad. Expression of StAR is not consistent with this trend, since its expression is similar between LC and brain. Nonetheless, our data collectively point to the woodpecker LC as a muscle that maintains a specialized capacity for *de novo* androgen synthesis.

### Constitutive vs. plastic regulation of enzyme expression

In a final analysis, we investigated whether muscular expression of androgen biosynthesis enzymes are linked to muscle use *per se* [40]. We therefore compared expression of these genes in the woodpecker LC between the breeding season and non-breeding season. Downy woodpeckers drum far more frequently during the breeding season, although they do produce occasional drums in the non-breeding season [36]. Thus, if expression of steroid biosynthesis enzymes were associated with display related muscle use, we would expect a strong seasonality in their expression level. Our data did not support this hypothesis, in that expression levels of all androgen biosynthesis genes were indistinguishable between the breeding and non-breeding seasons. We even confirmed that this assay for seasonal variation in gene expression was robust by comparing gonads from these same birds (Figure 3; Table 2). Indeed, when the testes of these individuals produce testosterone in the spring, all androgen biosynthesis genes are more abundantly expressed than during the winter when the gonads are regressed. These data are therefore consistent with a model in which muscle “activity” does not guide the expression of the steroid biosynthesis machinery in the myocyte.

**Table 2.** Summary of t-tests used to examine how steroidogenic ezyme expression changes in the testis, LC muscle, and PEC muscle of male downy woodpeckers across seasons

|  | **STAR** | **TPSO** | **Cyp11a1** | **3BHSD** | **Cyp17** | **17BHSD** | **5aR** | **AROM** |
| --- | --- | --- | --- | --- | --- | --- | --- | --- |
| **Testis** | t(5) = 4.29,<br>p < 0.01* | t(5) = 3.30,<br>p = 0.04* | t(5) = 6.28,<br>p < 0.01* | t(5) = 2.16,<br>p = 0.09 | t(5) = 7.27,<br>p < 0.01* | t(5) = 5.84,<br>p < 0.01* | t(5) = 2.86,<br>p = 0.05 | t(5) = 6.29,<br>p < 0.01 |
| **LC** | t(7) = -2.39,<br>p = 0.07 | t(7) =<br>-0.48, p = 0.65 | t(7) = 1.28, p =<br>0.25 | t(7) = 1.61, p =<br>0.17 | t(7) =<br>-0.28, p = 0.79 | t(7) = 0.83,<br>p = 0.44 | t(7) = 0.55,<br>p = 0.61 | t(7) = 5.53,<br>p < 0.01 * |
| **PEC** | t(7) =<br>-1.80, p = 0.13 | t(7) = 1.22,<br>p = 0.27 | n.d. | t(7) =<br>-1.32, p = .24 | t(7) = 0.52,<br>p = 0.63 | n.d. | t(7) = -0.45,<br>p = 0.67 | t(7) = 3.78,<br>p < 0.01 * |

\* denote significant differences after FDR correction for multiple tests n.d. denotes transcripts that were not detectable in a particular tissue

**Figure 3.**
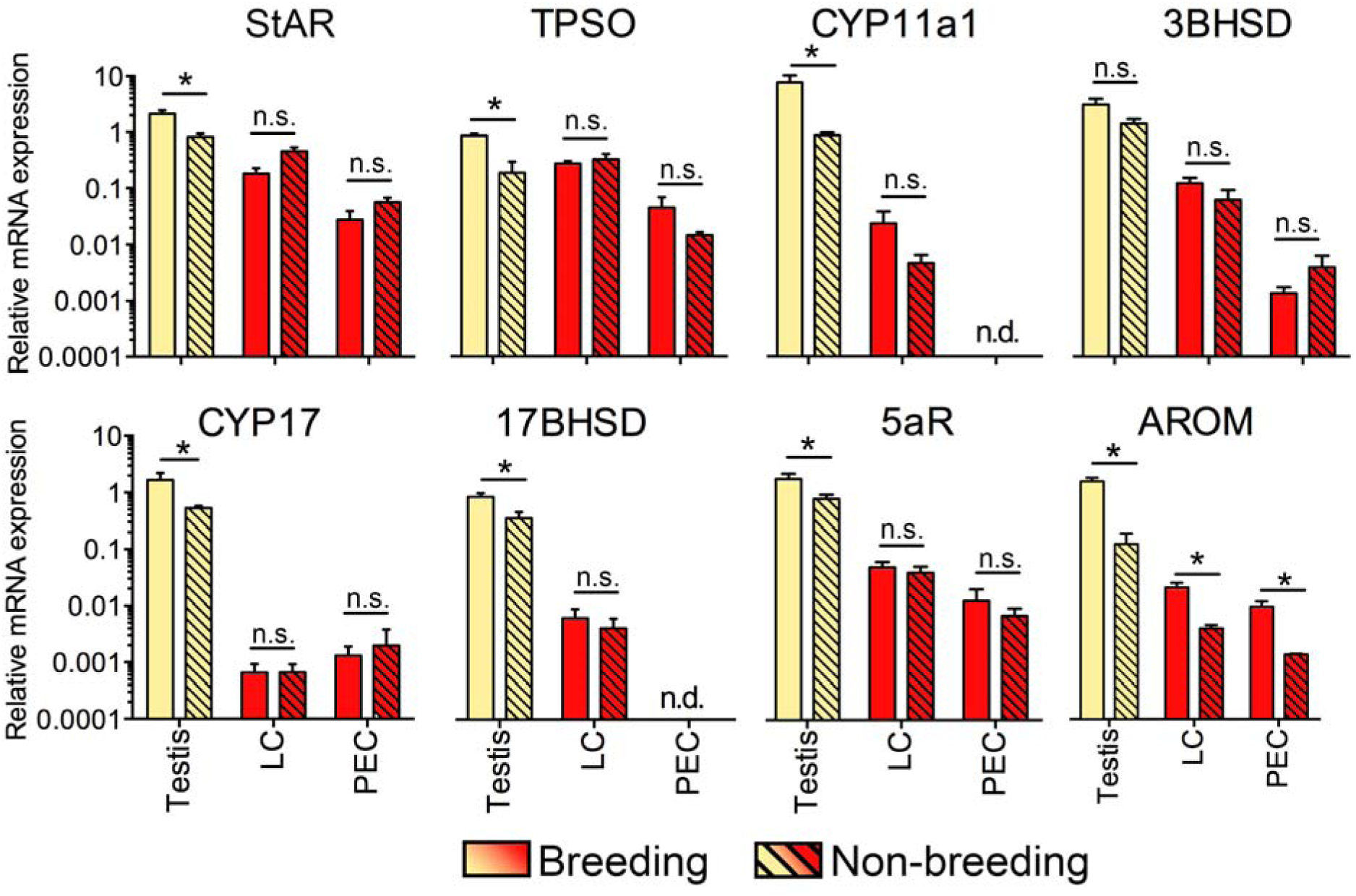
Expression of steroidogenic enzymes in the testis, *longus colli ventralis* (LC), and *pectoralis* (PEC) of breeding (solid bars) and non-breeding (hashed bars) male downy woodpeckers. Bars represent mean ±SEM. Asterisks denote significant differences (p <0.05) between the breeding and non-breeding season. n.s. denotes comparisons that did not significantly differ from each other.

## DISCUSSION

Androgenic steroids can act via skeletal muscle to mediate many elaborate social displays that animals use for courtship and competition [29, 30, 33]. Our current study uncovers an important new dimension to this process by showing that select muscles that govern such behavior are likely specialized for *de novo* androgen synthesis. This means that the muscle can make its own androgenic hormones, and thus allow steroidal regulation to occur outside the confines of the hypothalamic-pituitary-gonadal (HPG) axis. We provide several lines of evidence to support this idea. First, two different display muscles—the songbird SYR and the woodpecker LC—express all the transporters and enzymes necessary to convert cholesterol into bioactive androgens. This includes the expression of the genes that encode StAR and side-chain cleavage, which underlie the rate-limiting steps of steroid biosynthesis [37] and are typically absent in extra-gonadal tissue [25-27]. Second, expression levels of these genes are relatively greater in display muscles compared to a non-display muscle, and they even match levels in the brain (a known sites of steroid synthesis outside of the gonads). Lastly, the abundance of transcripts for genes involved in steroidogenesis are stable in display muscle across the year. This stands in contrast to the gonads, where transcript levels drop during the non-breeding seasons when circulating androgens are not made. Thus, the capacity of display muscle to make its own androgens is likely a constitutive property of the tissue itself, failing to change in response to fluctuations in circulating testosterone or behavior-related muscle activity [35].

### Functional significance

To our knowledge, our study is the first to indicate that skeletal muscle maintains the molecular machinery necessary for *de novo* androgen synthesis. Moreover, this study is also the first to suggest that this capacity is specialized specifically in muscles that actuate elaborate sexual displays. Thus, given the importance of androgen-muscle interactions to these behaviors, we hypothesize that myocytic synthesis of androgenic steroids supports fine motor control of movements that underlie effective communication. There are several ways in which these effects might occur. For example, androgens might act via the SYR to increase its muscle mass [11], enhance its capacity for neuromuscular transmission [41, 42], modulate its fiber type composition [43, 44], and/or calcium cycling machinery [45]. Such changes boost force production and the speed of muscle contraction, and thus facilitate rapid temporal features of song [44, 46]. Several of these acoustic parameters—including the on- and offset of vocalizations, rapid modulation of sound frequency and amplitude, as well as the rapid switching between contributions of the two sound generators—likely play a critical role in how a male’s singing ability is assessed by same- and opposite-sex conspecifics [47-49]. At the same time, it is possible that androgenic modulation of the rapid movement of the sound producing labia into the airstream lowers the required pressure for sound onset, which in turn leads to louder vocalizations [50] that enhance the potency of song [51, 52]. Past work certainly supports this view, showing that androgenic regulation of the SYR mediates several important acoustic elements of song, but not the motivation to produce it. For example, if androgen action in the SYR of canaries (*Serinus canaria*) is selectively blocked, then males produce songs with a disrupted syllable morphology and substantially reduced performance of ‘special trills’ [15]. This latter component to the display stimulates female copulation solicitation displays more than any other [53], suggesting that androgen-SYR interactions are vital to a male’s ability to copulate.

Many of these same principles also likely apply to the woodpecker LC. This muscle helps actuate the swift hammering behavior of the drum display [34, 54], meaning that the tissue must repeatedly contract and relax at an especially rapid rate [up to 20 Hz; 33]. Recent work shows that the LC is in fact specialized for this role, expressing high levels of genes that encode proteins to mediate quick calcium buffering and reuptake [55]. This should result in rapid muscle relaxation, which in turn increases contraction-relaxation cycling speeds. One way to maintain such machinery is through androgenic action, which regulates genes linked to calcium cycling in the myocyte [56, 57]. The woodpecker LC is likely no exception, given that it is heavily enriched with androgen receptor [35]. Thus, local synthesis of androgens in this muscle probably supports its ability to generate a fast drum, which is more effective during territorial combat [33]. Precedent for this idea comes from experimental work in a species of tropical bird, called the golden-collared manakin (*Manacus vitellinus*). In this species, androgens act on the wing musculature to dramatically increase the speed at which males perform acrobatic courtship displays [14, 58]. Females preferentially mate with faster displaying males, meaning that sexual selection for rapid display behavior likely drives the evolution of androgenic regulation of the wing muscles [59].

In light of our hypotheses about the function of *de novo* androgen synthesis in display muscle, we must also point out that these tissues express steroidogenic gene transcripts at levels roughly 1-5% of those in the gonads. These levels may at first seem low; however, we find that they are similar to those found in the brain, which is a known site of steroid synthesis [22, 60, 61]. Moreover, evidence suggests that cardiomyocytes express similarly low levels of steroidogenic enzymes relative to the testes [62], yet the cells are still fully capable of producing functional steroids *de novo* [24, 63]. It is therefore likely that display muscle—despite expressing low levels of the genes involved in steroidogenesis—can produce fully functional androgens.

### Evolutionary Significance

Why might display musculature evolve the capacity to synthesize androgens in the first place? One reason might be that it helps individuals temporally liberate their display behavior from functional barriers set by the hypothalamic-pituitary-gonadal axis. Most seasonally breeding individuals increase levels of circulating androgens at the onset of the reproductive season, particularly when males compete for resources and/or mates [64]. Thus, if this was the primary source of androgens to mediate display behavior by way of the muscle, then individuals could only display during these times of the year. Yet it is well established that several species readily perform socio-sexual signals outside of this narrow context — indeed, this ability is often adaptive [7, 8, 19, 65]. This is no exception for the species we examine herein, as all three sing or drum when circulating levels of androgens are otherwise basal [5, 30, 36]. Thus, the ability of the SYR and LC to make their own androgens likely allows individuals of each species to maintain the peripheral components of their display motor systems year-round. In this way, they would be capable of producing effective signals regardless of androgen levels in the blood, as long as the brain could still initiate such behavior (see below).

Androgen synthesis in display muscle may also help mitigate fitness ‘costs’ normally associated with these hormones. Indeed, gonadal androgens suppress immune function, diminish fat reserves, and slow the process of wound healing [66-68]. Prolonged exposure to circulating androgens outside periods of territoriality and/or breeding also interferes with parental care [69]. All these effects negatively impact viability and reproductive success [70, 71], and thus selection favors mechanisms that reduce or offset their impact. As such, low-levels of androgenic synthesis in the target muscles likely provides a solution to this ‘evolutionary problem’ by allowing the performance of the androgen-dependent behavior without high levels of circulating androgens [72].

### A new dimension to androgen-muscle relationships

Our data stress the potential importance of androgenic regulation of muscle involved in complex animal behavior, a view currently gaining acceptance [73]. However, these findings should not detract from the long-held view that circulating androgens are critical to the organization and activation of many sexual behaviors. Both of these models likely apply to most vertebrates, and it is the integration of these two viewpoints that forwards our thinking about how steroid systems accommodate reproductive traits. Accordingly, we hypothesize that circulating brain-level androgen action helps regulate facets of reproductive behavior related to sensory integration, arousal, motivation, and/or attention [74]. By contrast, we expect that androgens act through muscle to mediate performance-related aspects of behavior, such as agility, speed, endurance, etc. [75]. In this way, both “top down” and “bottom up” channels of steroid action are necessary to fully actuate certain behavioral traits. Our current findings therefore suggest that such channels are not necessarily coupled to the same hormonal source (i.e., HPG axis), rendering the hormonal mechanisms far more labile—both mechanistically and evolutionarily—than previously thought.

### Conclusions

Here, we find that important display muscles in three different avian species maintain the machinery needed to undergo *de novo* androgen synthesis. This trait appears to be a constitutive specialization of the muscle itself, resembling a similar specialization of the brain for the ability to produce its own steroid. We hypothesize that androgen synthesis in display muscle is essential for maintaining its ability to actuate necessary signaling behavior outside of the breeding season, and thus is an adaptation to enhance social communication behavior.

## METHODS

### Animals

All appropriate federal, state, and university authorities approved of the methods described below. During March and April, we passively captured male downy woodpeckers (*n*=4) and white-breasted nuthatches (*n*=3) using mist nets in the woodlands of Forsyth County, North Carolina (USA). Individuals were immediately euthanized via cervical dislocation to keep the LC muscle intact and flash frozen on dry ice. We stored specimens at −80°C until they were later processed, at which point we dissected out the whole brain, gonads, SYR muscle (nuthatches only), LC muscle, and PEC muscle. During these dissections, we also verified that the gonads from each individual were enlarged to a size consistent with an actively breeding bird. During November to December of the same year, we passively captured and dissected a set of non-breeding down woodpeckers (*n*=3) using the same techniques described above. We similarly verified that the testes were regressed to a size consistent with a non-breeding bird.

Finally, we euthanized adult (age >120 days) male zebra finches (n=5) with an overdose of isoflurane before dissection. The birds had been individually housed in a 31.8 cm × 22.9 cm × 27.9 cm wire cages with newspaper lining. They were fed a mixture of red and white millet, canary seed, and water *ad libitum*, and they were supplemented with peas and corn every other day.

### RNA extraction and reverse transcription

We homogenized each tissue sample in TRIzol Reagent™ using a rotor/stator homogenizer set to medium speed. We extracted total RNA from these samples with a Zymo Direct-zol RNA miniprep kit (Zymo Research, Irvine, CA), in which we included an initial phenol-chloroform separation of RNA as per manufacturer instructions. Next, we treated RNA samples with DNAse and reverse transcribed 1 μg of RNA, using SuperScript IV Reverse Transcriptase (Invitrogen). Each reaction occurred for 10 min at 55°C, followed by 10 min at 80°C.

### PCR and Sequencing

Using cDNA from the extracted tissues, we amplified the different genes that encode the proteins that mediate steroid biosynthesis (Figure S1). The primers for the initial PCR reactions were designed from either the downy woodpecker or the zebra finch genomes to conserved regions of each gene (Table S1). All PCR reactions contained 40 ng of cDNA, 0.5 μM of forward primer, 0.5 μM of reverse primer, OneTaq 2x Mastermix (New England Biology). Reactions were run at 96 °C for 5 min, followed by 40 cycles of 96°C for 30 s; 57–60 °C for 30 s; and 68°C for 30 s. Each reaction was completed with a final extension step at 68°C for 5 min.

Resulting PCR products were imaged on a 1% agarose gel to verify that fragments matched their expected size. We then purified the PCR products using a GeneJet PCR purification kit (ThermoFisher) and had them sequenced (Eton Bioscience). For cyp11a1, STAR, and 17BHSD, resulting purified PCR products were amplified a second time using the parameters described above to generate a high enough concentration of DNA for accurate and efficient sequencing. All PCR products matched the expected gene of interest.

### Quantitative PCR (qPCR)

We used quantitative real-time PCR (qPCR) to measure each steroidogenic enzyme’s gene expression in the different tissues of our three species relative to GAPDH (housekeeping control). In all three species, we confirmed that GAPDH was acting as an appropriate endogenous control by verifying that there were no significant differences in between tissues in GAPDH expression (all p values >0.2). For this analysis, reactions were run in an Applied Biosystems QuantStudio3 machine. We used the gene sequences obtained from our PCR runs to generate species-specific primers (Table S2), and we ran each reaction with 100 ng of cDNA, 0.9 mM of each primer, and SYBR Green Master Mix (Applied Biosystems). For all genes, we included negative controls that consisted of all elements in the reactions described above except cDNA. Reactions were run using the following parameters: 50 °C for 2 min, 95 °C for 10 min, followed by 40 cycles of 95 °C for 15 s, and 60 °C for 1 min. We added a final dissociation stage to the end of the reaction process, which consisted of 95 °C for 15 s, 60 °C for 30 s, and finally 95 °C for 15 s. All negative controls were undetectable.

All measurable samples were detectable with CT values below 35. However, in a handful of cases, mRNA for a given gene interest was not detectable after 40 cycles (see Table S3d)—we therefore list count these tissues as not expression the gene of interest [i.e., ‘not detectable’ (n.d.); See Figure 3]. Reaction efficiencies were between 90% and 110%, and we used the standard curve method to measure relative expression (i.e., quantity of gene of interest /quantity GAPDH). The standard curves used in all the current experiments were generated serially diluted (1:4) pooled cDNA.

### Data analysis

To next investigate whether there are tissue differences in mRNA expression of each of the steroidogenic enzymes, we performed linear mixed models (LMM) for each species with tissue as a fixed factor. To account for the non-independence of gene expression levels between tissues in the same animal, we used individual identity as a random factor in each model. In all analyses, significant main effects were further examined with *post-hoc* comparisons, using FDR corrections to account for multiple contrasts. We used independent t-tests to determine how the capacity for steroid synthesis changed in the testis, LC muscle, and PEC muscle across seasons in downy woodpeckers. Furthermore, we performed FDR corrections to account for the number of LMMs and t-tests performed. All analyses were performed in R (v3.3.2).

## Data accessibility

The data will be uploaded to Dryad upon acceptance of the manuscript.

## Ethics

All the experiments in this study were conducted with approval from the Wake Forest University Institution Animal Care and Use Committee (# A16-187).

## Author contributions

ERS participated in study design, collected the animals, performed lab experiments, analyzed data, and wrote the manuscript. FG and MJF participated in study design, analyzed data, and wrote the manuscript.

## Acknowledgements

We thank Johnny Petersen, Emma Vivlamore, and Ryan Walsh for field and laboratory assistance.

